# Lung Bronchial Epithelial Cells are HIV Targets for Proviral Genomic Integration

**DOI:** 10.1101/2020.06.01.126821

**Authors:** Dinesh Devadoss, Shashi P. Singh, Arpan Acharya, Kieu Chinh Do, Palsamy Periyasamy, Marko Manevski, Neerad Mishra, Carmen Tellez, Sundaram Ramakrishnan, Steve Belinsky, Siddappa Byrareddy, Shilpa Buch, Hitendra S. Chand, Mohan Sopori

**Author notes:** Equal Contributions (Dinesh Devadoss, Shashi P. Singh). **Corresponding Authors:** Mohan Sopori, Ph.D., Lovelace Respiratory Research Institute, Albuquerque, NM 87108, USA, Hitendra S Chand, Ph.D., Herbert Wertheim College of Medicine, Florida International University, Miami, FL 33199, USA.

## Abstract

In the era of highly active anti-retroviral therapy (HAART), obstructive lung diseases (OLDs) are common among the people living with HIV (PLWH); however, the mechanism by which HIV induces OLDs is unclear. Although human bronchial epithelial cells (HBECs) express HIV coreceptors and are critical in regulating lung immune responses, their role in transmitting HIV remains unclear. Herein, we present evidence that HIV-1 infects normal HBECs and the viral DNA is integrated in the genome to establish the viral latency. To prove that HIV productively infects HBECs, we demonstrate: (a) along with CXCR4, HBECs express the HIV-receptor CD4, and are induced to express CCR5 by IL-13 treatment; (b) following infection with HIV, HBECs produce HIV-p24 and contain the latent HIV provirus, which is activated by endotoxin and/or vorinostat; (c) DNA from HIV-1 infected HBECs contains the HIV-specific *gag* and *nef* genes, along with *Alu* sequences, confirming the integration of HIV in the host DNA; (d) the lung epithelial cells of HIV-infected subjects and SHIV-infected cynomolgus macaques are positive for HIV-specific transcripts. Thus, these studies suggest that HIV establishes latency in lung epithelial cells, making them potential HIV reservoirs. The long-living lung epithelial cells, activated by commonly encountered lung infections, might represent an ideal HIV target/reservoir, contributing to OLDs and other HIV-associated lung comorbidities.

## INTRODUCTION

The combined antiretroviral therapy (cART) or HAART has improved the life span of people living with HIV (PLWH), however, the major hurdle in achieving complete cure of HIV is the persistence of latent reservoirs i.e. the cells harboring quiescent HIV provirus (1–3). Resting CD4+ T cells are identified as one of major latent reservoirs; however, HIV infections are anatomically found in all vital organs including lungs (4, 5), but the presence of latent non-immune cell HIV reservoirs in these tissues remains unclear. These putative reservoirs may contribute significantly to the comorbidities of the organ. Frequency and the early onset of OLDs such as chronic bronchitis (CB) and chronic obstructive pulmonary disease (COPD) are significantly higher among PLWH (6–9). Airway mucus hypersecretion is associated with chronic bronchitis and COPD (10) and we have shown that, even after cART, lungs from HIV-infected humans and SIV-infected macaques contain large amounts of mucus (11). In the lung, the mucus is produced by specialized secretory lung epithelial cells (goblet cells), which are also the key effectors of chronic bronchitis and COPD (12). We have demonstrated that HIV-gp120 stimulates mucus formation in normal human bronchial epithelial (NHBE) cells (11) and epidemiological evidences suggest that HIV is an independent risk factor for the development of COPD (13, 14).

The role of epithelial cells as one of the cellular targets of HIV infection is not clearly resolved. Foreskin epithelial cells express HIV coreceptors (15) and when the cells are infected with herpes simplex virus 2, the cells internalize HIV virions (16). Although the virions do not replicate, they can be transferred to lymphocytes by coculturing (17). On the other hand, NHBE cells express CXCR4 (C-X-C motif chemokine receptor-4) and the X4-tropic gp120 induces mucus in these cells (11). Similarly, X4-tropic, but not R5-tropic, HIV impairs epithelial barrier integrity in NHBE cells (18). Also, cynomolgus monkeys (CMs) infected with simian-adapted HIV (SHIV) contain large number of mucus containing goblet cells and a significant percentage of SHIV-infected airway epithelial cells are HIV-gp120-positive (19); however, whether HIV productively infects airway epithelial cells is still debatable (18, 20). This question can be answered definitively, if the HIV-infected bronchial epithelial cells are shown to contain activatable integrated HIV provirus. In this study, we demonstrate that, following HIV infection, NHBE cells produce HIV p24 and become carriers of latent HIV provirus that can be activated by treating the virus-containing cells with lipopolysaccharide (LPS) and/or vorinostat. Moreover, DNA from HIV-1 infected NHBE cells contains HIV-specific genes and HIV- and SHIV-infected lung epithelial cells express HIV-specific RNAs.

## RESULTS & DISCUSSION

### Human bronchial epithelial cells intrinsically express CD4 and CXCR4 *in-vitro*, and can be induced to express CCR5 HIV Co-receptor

Besides CD4, the CXCR4 and CCR5 (C-C motif chemokine receptor-5) are known HIV co-receptors (21). The expression of CXCR4 on NHBE cells has been amply documented (11, 22, 23), and we have also previously demonstrated that following infection, NHBE cells produce mucus in response to X4-but not to R5-tropic HIV-gp120 (11); however, the expression CCR5 on these cells is not unequivocal (18, 20). During carcinogenesis, epithelial cells have been shown to express CCR5 that enhances their resistance to cytotoxicity (24). We first analyzed the expression pattern of CD4 and other HIV co-receptors in normal disease-free NHBE cells. Based on the qPCR and immunoblot analysis, NHBE cells expressed CD4 that was more than fifty-fold higher at mRNA levels (**Fig. 1A**) and around five-fold higher at protein levels (**Fig. 1B**), respectively, in comparison with A549 cells (a human lung adenocarcinoma cell line of lung epithelial origin, used for experimental reference). These NHBE cells also showed high immunopositivity for CD4 (**Fig. 1C**) and CXCR4 (**Fig. 1D**) with 3-fold more abundance of CD4 compared to CXCR4 levels (**Fig. 1E**). On the other hand, CCR5 was expressed at very low levels in naïve NHBE cells cultured in submerged conditions; however, these cells showed an induced expression of CCR5 following 48 h stimulation with a Th2 cytokine, IL-13 at 1 ng/ml (**Suppl Fig. S1A**). The IL-13-induced CCR5 expression (**Suppl Figs. S1B and S1C**) correlated with susceptibility of NHBEs to R5-tropic HIV-1 (HIV_BaL_), leading to higher HIV-gp120 positivity (**Suppl Fig. S1D**) in IL-13 treated cells compared to the non-treated (NT) controls.

**Figure. 1:**
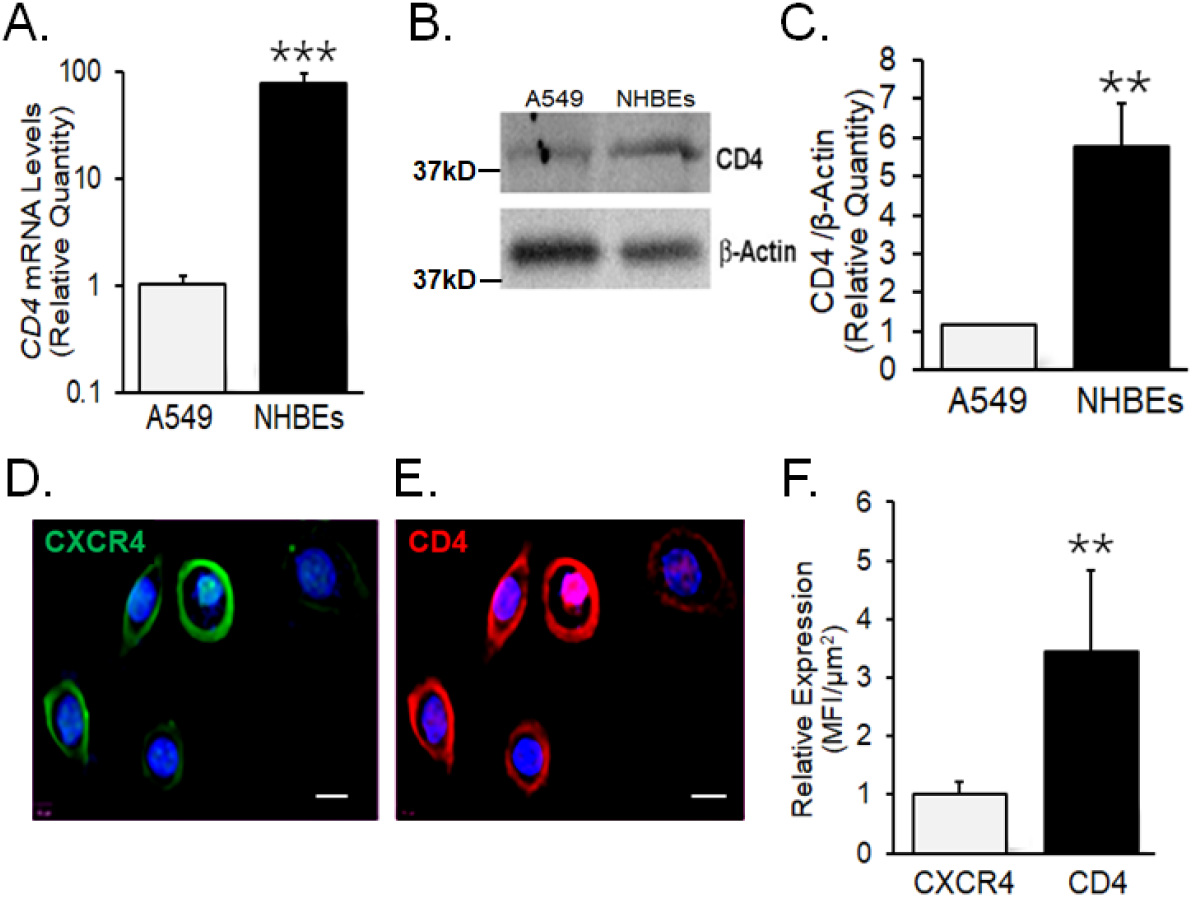
Human airway epithelial cells express HIV receptor and co-receptors. **(A.)** qPCR analysis of CD4 mRNA levels in NHBEs compared to A549 cells. **(B.)** Western blot analysis of CD4 protein levels in NHBE and A549 cells with β-actin as loading control. **(C.)** Densitometric analysis of CD4 protein levels compared to A549 cells. Representative micrographs of NHBE cells showing **(D.)** CXCR4 (green) and **(E.)** CD4 (red) immunopositivity along with DAPI-stained nuclei (blue); scale −5μ. **(F.)** Relative expression of CXCR4 and CD4 in NHBEs as assessed by measuring the Mean Fluorescence Intensity (MFI) per unit area. Data shown as mean±SEM; n=3/gp; **p<0.01; ***p<0.001

### HIV co-receptors are expressed *in-vivo* in lung epithelium of nonhuman primates

Next, we analyzed the CD4 and CXCR4 receptor expression in the lung epithelium of nonhuman primates. Archived lung tissues from the cynomologus macaques (CMs) that were exposed to CS for 27 weeks and/or infected with SHIV at 11^th^ week to obtain 4 experimental groups: fresh air control (FA), CS-exposed, SHIV-infected, and CS+SHIV, as described recently (19). All SHIV-infected macaques received daily injections of cART (Tenofovir and Emtrictabine) starting at 2 weeks post infection until euthanasia. We analyzed the expression of HIV secretory protein, transactivator protein (Tat) which regulates HIV transcription inside the host. There was very high expression of immunoreactive Tat was observed in lung epithelial cells of SHIV-infected CMs (**Fig. 2A**) that have been successfully treated with cART, and the expression was significantly higher (more than four-fold) in the lungs of CS-exposed and SHIV-infected (CS+SHIV) macaques (**Fig. 2B**). Further immunofluorescence analyses showed that, compared to control, the airway epithelial cells (pan-cytokeratin or pCK-positive cells) from the CS-exposed group had significantly upregulated expression of CD4 (**Fig. 2C**) and CXCR4 (**Fig. 2D**) in CS+SHIV macaques, supporting our in-vitro findings in human NHBEs (**Fig. 1**). The expression of HIV co-receptor, CCR5 (**Suppl. Fig. S2**) was also confirmed in-vivo in the CM lung epithelium.

**Figure 2.**
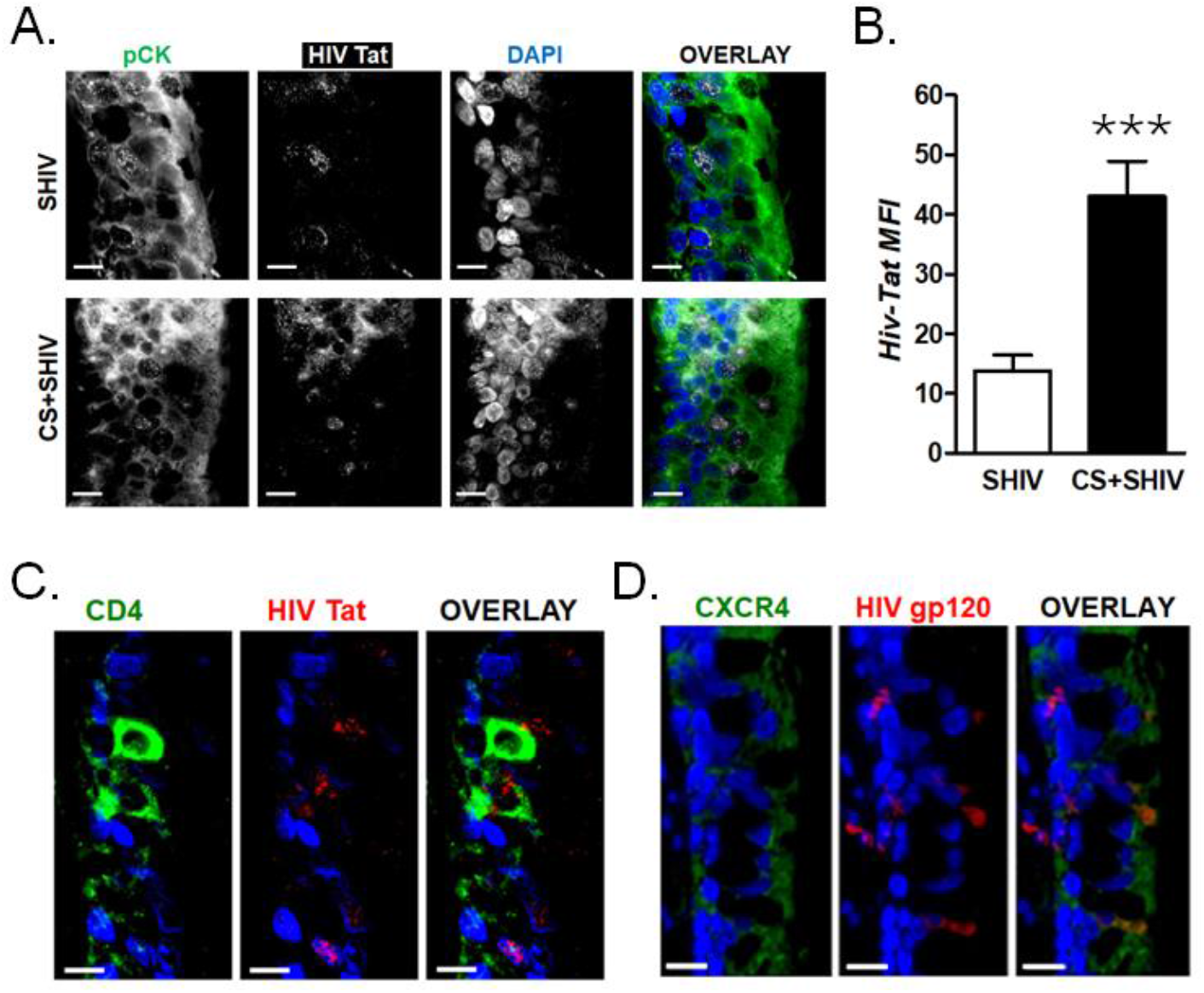
CS exposure induces CD4 and CXCR4 expression in the lung epithelial cells of SHIV-infected nonhuman primates. Archived formalin-fixed and paraffin-embedded (FFPE) lung tissue sections (5 μm) from CM exposed to CS or CS+SHIV were used for the immunofluorescence analysis. (**A**) Representative micrographs of bronchial airways of SHIV-infected CMs exposed to CS showing immunopositivity for HIV-Tat (white), pan-cytokeratin (pCK, green), along with DAPI-stained nuclei (blue), scale - 5μ. (**B**) Mean Florescence Intensity of HIV-Tat. Data shown as mean±SEM; n=4/gp; ***p<0.001. Representative micrographs of bronchial airway epithelial cells showing (**C.**) CD4 (green) and (**D.**) CXCR4 (green) expression in CS+SHIV-infected CMs with the co-detection of HIV gp120 (red) and the DAPI-stained nuclei (blue); scale - 5μ.

### Blocking of HIV receptors or co-receptors significantly reduces HIV infectivity of 3D-cultured NHBE cells

To confirm the of HIV-1 uses classical HIV receptors and co-receptors to infect, 3D-cultured NHBE on air-liquid interface (ALI) were pre-incubated with anti-CD4 and/or anti-CXCR4 antibodies for 1 h at 37°C and then infected with CXCR4-tropic HIV-1_LAV_. At 24 h post-infection, cellular RNA was isolated and assayed for HIV-specific LTR RNA levels. As seen in Figure 3A, compared to control immunoglobulin-treated cells, anti-CD4, anti-CXCR4, or anti-CD4+anti-CXCR4 significantly reduced the HIV-RNA expression. However, the level of the mucin *MUC5AC* was more sensitive to anti-CXCR4 than anti-CD4, but the combination (anti-CD4+anti-CXCR4) potently reduced the viral RNA expression (**Fig. 3B**). We have previously shown that HIV-gp120-induced mucus formation and MUC5AC expression in NHBE cells is suppressed by blocking CXCR4 (11). Along with CD4, HIV coreceptors have been strongly implicated in optimal HIV-gp120 binding and viral infection (25). These results suggest that HIV-1 enters NHBE cells via CD4 and CXCR4, but the mucus formation primarily depends on the binding of the virus to the cell surface and may not necessarily require a productive viral infection.

**Figure 3.**
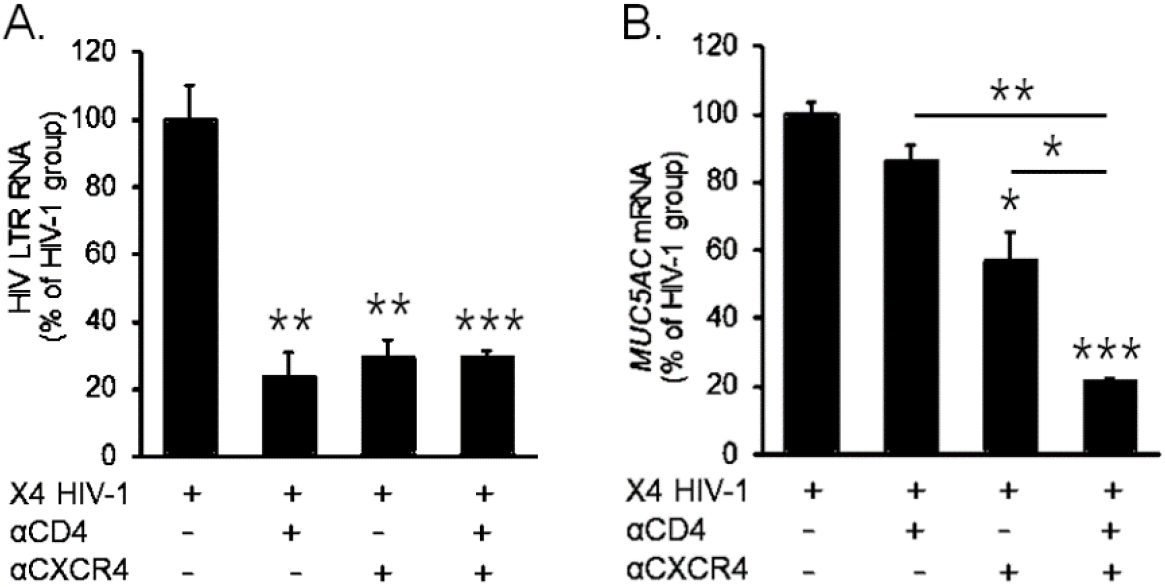
Blocking HIV-coreceptors inhibits HIV infectivity in NHBE cells. Relative quantities of HIV-1 LTR RNA levels (**A.**) and *MUC5AC* mRNA (**B.**) in NHBE cells pre-treated with either anti-CD4 or anti-CXCR4 antibodies followed by infection with X4-tropic HIV_LAV_ and 48 h post-infection cells were harvested and analyzed. Data shown as mean±SEM; n=3/gp; *p<0.05; **p<0.01; ***p<0.001

### X4-tropic HIV-1_LAV_ infects differentiated NHBE cells and establishes latency

To determine whether HIV infects and establishes latency in differentiated NHBE cells, NHBE cells were grown on ALI and infected with HIV-1_LAV_ either apically or basolaterally in trans-well cell culture. After 2 h post-infection, media was removed, and the trans-well filters were washed on both sides 4 times with warm ALI culture media and each wash was tested for HIV-p24 to confirm the removal of input virus. As shown in **Figure 4A** there was no detectable p24 in the 4^th^ wash and was considered as 0 h time point. Cells were incubated in fresh medium and, at the indicated times, media aliquots were collected from both apical and basolateral sides and assayed for HIV p24. Cells were infected with basolateral side, The level of p24 were increased in the cultures infected basolaterally reaching maximum (30 pg/ml) at 1 day (24 h) post-infection; thereafter, the p24 levels started declining, reaching very low levels by day 7 (**Fig. 4A**). On the other hand, the p24 levels of the apically infected cultures remained undetectable throughout this period (data not shown). These results suggest that the airway epithelial cells are infected by HIV via the basolateral but not the apical side and may partly contribute to the lack of HIV transmission via the oral route.

**Figure 4.**
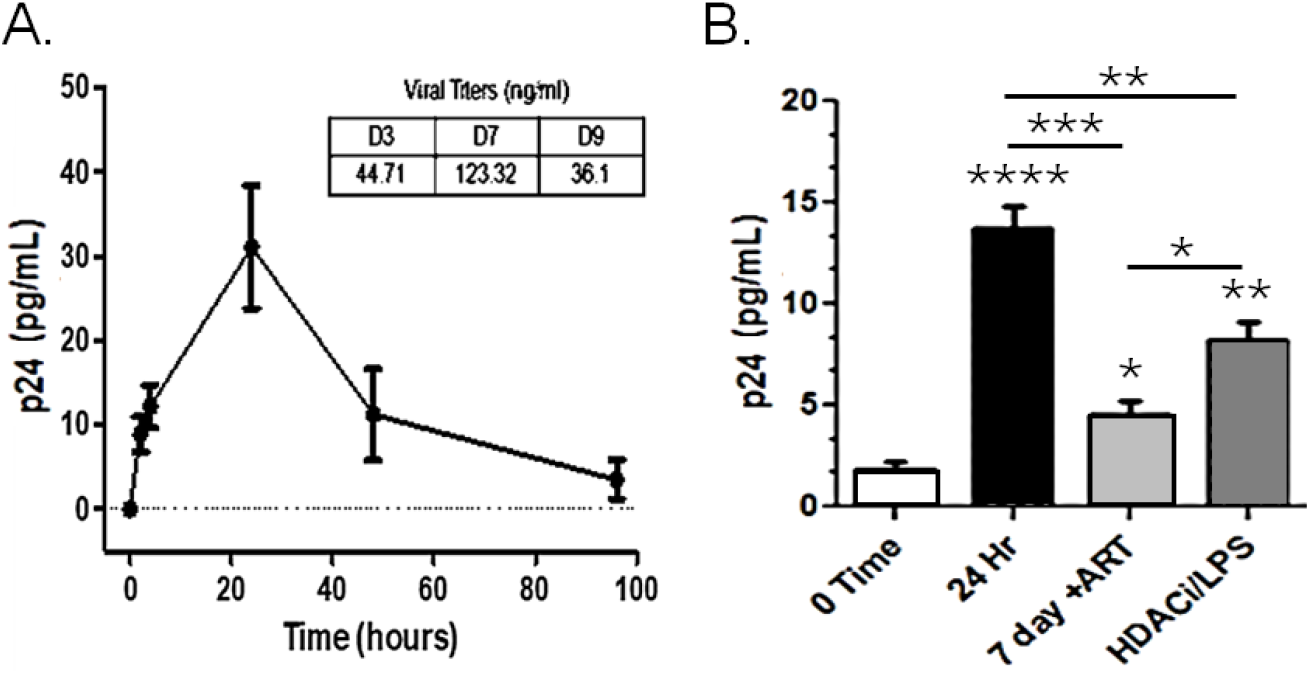
Differentiated NHBE cells were infected by X4-tropic HIV-1 and act as HIV-1 reservoir. (**A.**) Expression kinetics of p24 (pg/ml) measured in cell culture media collected from the bottom of trans well after X4-tropic HIV-1(LAV) infection; inset-viral load (ng/ml) of day 3, 7 and 9. (**B.**) HIV-1 p24 levels after vorinostat and LPS treatment of HIV-1 infected cells. Data shown as mean±SEM; n=3/gp; *p<0.05; **p<0.01; ***p<0.001; ***p<0.0001

In tissue reservoirs, HIV is known to establish latency and, in cell cultures, the virus is activated by various reactivation stimuli, including histone deacetylase (HDAC) inhibitors (26, 27) and several toll-like receptor (TLR) agonists like LPS (28–30). To determine if HIV infection of NHBE cells establishes viral latency in the surviving cells from 7-day post-infection Transwell cultures, the cells were treated with the latency reversing agents, LPS (1 μg/ml) and/or the HDAC inhibitor, vorinostat (300 nM) that is shown to activate the HIV provirus in vivo and in vitro (31, 32). At 24 h after the treatment, the HIV-1 Gag p24 levels were significantly upregulated (**Fig. 4B**). This accords well with the observation that vorinostat treatment of HIV provirus containing T cells from patients breaks their latency, leading to increased HIV-specific RNA in the cell (27). Activation of HIV provirus is invariably associated with cell apoptosis (33) and, indeed, following each of these treatments, essentially all the cells on the Transwell filters were dead (data not shown). These observations clearly show that HIV infects and establishes latency in differentiated NHBE cells. Besides, the respiratory tract is constantly exposed to viruses and bacteria and many of these activate TLRs on the cells. Therefore, it is feasible that respiratory infections will promote activation of HIV proviruses that are integrated within the airway epithelial cells, making the lung epithelial cells as an important reservoir of HIV.

### HIV-1 DNA is integrated in the NHBE cell genome

For a productive HIV infection, the reverse transcribed HIV DNA enters the nucleus to integrate into the cellular DNA and this integration of the provirus into the host genome is a central event in HIV pathogenesis (34). HIV proviruses integrate at many sites in the host genome (35) and, in resting cells, the integrated viral genome may remain essentially silent, leading to latency and viral replication primarily through clonal expansion (36). Therefore, to identify a HIV target, it is imperative to show that the viral DNA is integrated into the cellular DNA. To demonstrate this, we focused on genes that follow the 5’-LTR and precede the 3’-LTR of HIV (i.e., gag and Nef genes, respectively). We amplified these HIV genes from the DNA isolated from control (non-infected) and HIV-1-infected differentiated NHBEs and sequenced the amplified products. Briefly, DNA was isolated from control and HIV-infected NHBE cells at 24h post-infection. Two approaches were used to identify viral genes in the host cell DNA.

In the first approach, we amplified HIV *gag* by nested PCR. A single band at 1.52 Kb was present on the 1.0% agarose gel that was absent in the control DNA (**Fig. 5A**). The product was extracted from the gel and sequenced. The sequence was >99% identical to human HIV-1 *gag* by blast analysis using both forward primers (**Fig. 5B**) and reverse primers (**Suppl. Fig. S2A**). Similarly, Nef gene was amplified by nested PCR and isolated by gel electrophoresis. A single band on the gene at 730 bp (**Suppl. Fig. S2B**) was sequenced and showed >99%identity with published HIV-1 *Nef* sequence. The primers for the nested PCR and the sequence are provided in the online supplemental data (**Suppl. Fig. S2C**).

**Figure. 5:**
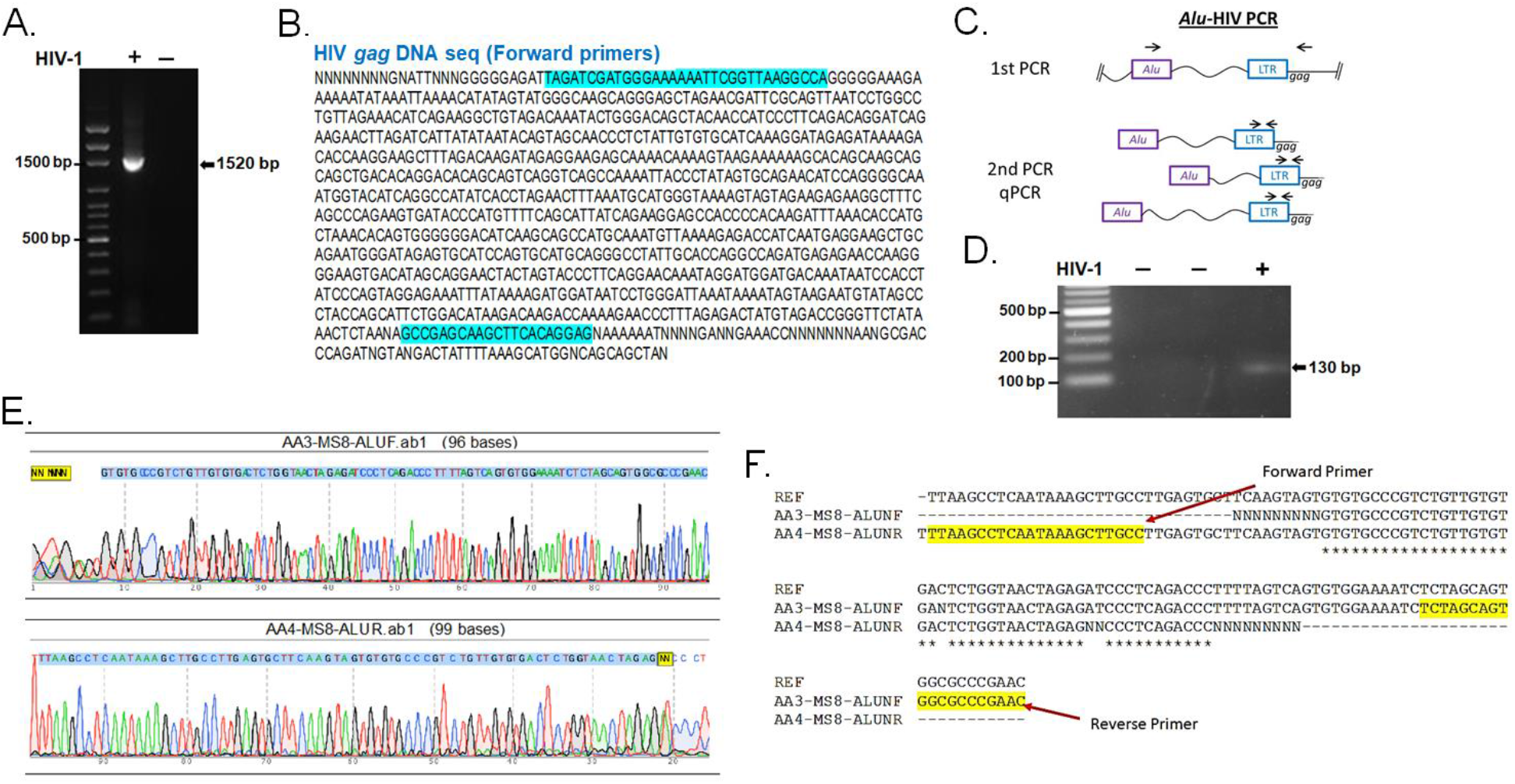
Amplification of HIV gag DNA and Alu-PCR analysis to confirm the integration of HIV-1 proviral genome in NHBEs. **(A.)** Agarose gel analysis of the HIV gag PCR amplicon (1520 bp) amplified from the DNA isolated from HIV-infected NHBEs. **(B.)** HIV gag DNA sequence of the amplicon obtained using the forward primers and 5’-LTR sequence of HIV are highlighted in blue color. **(C.)** Schematics of the nested-PCR approach used to confirm the HIV DNA integration in infected NHBEs. **(D.)** Agarose gel analysis of the nested PCR amplicon showing amplification of 130 bp band from 5’ LTR region of HIV-1. (**E.**) Electropherogram of the 130 bp PCR amplicon generated using Forward and Reverse primers confirms the presence of HIV proviral genome (The 5’-LTR sequence of HIV is indicated with blue shade). **(F.)** BLAST sequence analysis of HIV Nef DNA sequence obtained from sequencing of the 703 bp PCR product. The primer sequences used are highlighted in yellow.

In the second approach, we employed a two-step, *Alu-gag* PCR assay to confirm the presence of integrated HIV-1 proviral DNA. This was done by isolating DNA from uninfected controls and HIV-1 infected NHBE cells and subjected to nested PCR approach as described schematically (**Fig. 5C**), Briefly, the 1^st^ round PCR was performed with forward primer that anneals to Alu repeat elements in the human genome and the reverse primer anneals to HIV-1 *gag* gene. In the second round of PCR, the 1^st^ PCR amplicon was used as a template to amplify a 130 bp region of the 5’-LTR of HIV-1 genome as identified by agarose gel analysis (**Fig. 5D**). Finally, DNA sequencing was performed to confirm that 130 bp amplicon generated from the 5’ LTR region of HIV-1contains intact sequences of LTR-*gag* (**Fig. 5E**). These results clearly show that host DNA isolated from the HIV-infected NHBE cells contains integrated HIV proviral DNA. Presence of the 130 bp PCR amplicon in the DNA sample extracted from HIV infected cells, confirm the presence of integrated Proviral DNA in the NHBE genome. The PCR amplicon was sequenced with amplification primers to confirm target sequence. The sequence data exactly match with reference sequence (>K02013.1 HIV-1 1, isolate BRU, complete genome (LAV-1). The Forward Primer sequence detected in the reverse complement sequence generated using reverse primer and the Reverse Primer sequence detected in the sequence generated using forward primer (**Fig. 5F**), confirming the genomic integration of HIV provirus in NHBEs.

### HIV RNA is present in lung epithelial cells from SIHV-infected macaque and HIV-infected human lungs

We have previously demonstrated that HIV- and SIV-infected lungs from humans and monkeys, respectively, contain HIV-gp120 immunoreactive cells (11). More recently, we have shown that a significant number of airway and alveolar epithelial cells in SHIV-infected CMs are HIV-gp120-positive and the number of these cells are essentially doubled in SHIV+CS-exposed lungs (19). To determine that HIV replicates in airway epithelial cells in-vivo, we performed RNA FISH by RNAScope^®^ technology (Advanced Cell Diagnostics, Biotechne Inc.) using commercially available HIVgag-pol probes (ACD, Bioteche Inc.). As seen in **Fig. 6A**, HIV-specific RNA is present in SHIV-infected pan-cytokeratin (pCK) positive lung epithelial cells and this expression was three-fold higher in the animals, which were also exposed to CS i.e. CS+SHIV group (**Fig. 6B**). Similarly, unlike the uninfected controls, the archived airway sections of HIV-infected human subjects contain significant amount of HIV-specific RNA (20-fold higher than uninfected) and the RNA persisted (14-fold higher than uninfected) in HIV subjects on HAART (HIV+HARRT group) (**Figs. 6E** and **6F**). These results suggest that lung epithelial cells in humans and macaques are targets of HIV infection.

**Figure. 6:**
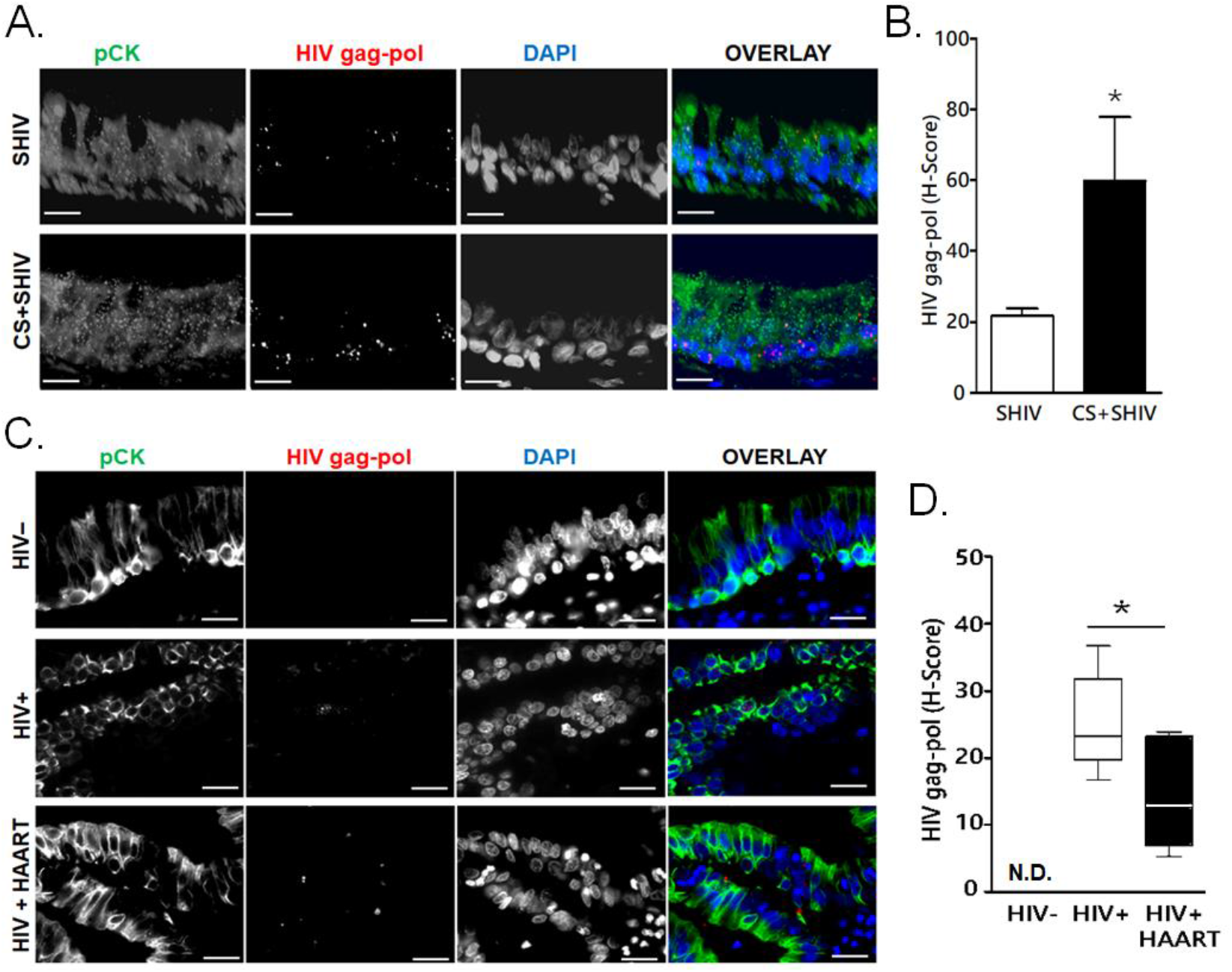
Expression of HIVgag-pol RNAs in the lung epithelial cells of SHIV-infected macaques and HIV-infected human subjects. Archived 5 μm FFPE lung tissue sections were used for HIVgag-pol RNA detection using RNAscope and immunostained for the epithelial cell marker, pan-cytokeratin (pCK). (**A.**) Representative micrographs of bronchial epithelial cells showing pan-cytokeratin, pCK (green) and HIVgag-pol (red) colocalization along with the DAPI-stained nuclei (blue) from SHIV- and CS+SHIV-infected groups; Scale - 5μ. (**B.**) Quantification of HIVgag-pol expression (H-score); *p<0.05. (**C.**) Representative micrographs of bronchial epithelial cells showing pCK (green) and HIVgag-pol (red) colocalization along with the DAPI-stained nuclei (blue) in archived lung tissues from HIV uninfected (HIV-), HIV-infected (HIV+), and HIV+HAART-treated subjects. Scale - 5μ. (**D.**) Quantification of HIVgag-pol expression (H-score). N.D. - Not detected. Data shown as mean±SEM; n=4/gp; *p<0.05.

In this study, we report that differentiated NHBE cells normally express receptors or co-receptors CD4 and CXCR4, and when exposed to X4-tropic HIV, the cells produce HIV p24. Moreover, a chronic infection of NHBE cells induces latent HIV proviral infection and the provirus can be activated by latency activating agents such as LPS and/or vorinostat that leads to cell death. can; however, upon treatment with IL-13, the cells also express CCR5. When short time/acute exposure of X4-tropic HIV, NHBE cells produced HIV-p24 and become carriers of latent HIV provirus which confirmed with reactivation by LPS and/or vorinostat along with apoptotic analysis. Further, Alu PCR confirmed the presence of integrated proviral DNA in HIV-infected NHBE cells, and the FISH analysis of lung epithelial cells of HIV-infected subjects were positive for HIV specific RNA. In the same way, CS exposed SHIV-infected NHPs express CD4 and HIV co-receptors, HIV-specific RNA and other viral counterparts like gp120 and HIV-Tat even after ART. In addition, blocking of HIV co-receptors inhibit HIV-infectivity and reduce mucus production in NHBE cells, which indicates HIV infects NHBE cells via the entry mediated by CD4 and other co-receptors.

Lung bronchial epithelial cells are crucial innate immune cells that actively help eliminate various environmental insults and pathogens. Other than tissue resident/infiltrate lymphoid and non-lymphoid immune cells, many epithelial cells express HIV receptor CD4 and co-receptors CXCR4 and CCR5 enabling either productive or non-productive infections (37–39). Identification of all host tissue HIV reservoirs is the first step towards complete eradication of HIV virions in the affected people. The respiratory tract is constantly exposed to airborne viruses and bacteria that can engage TLRs on the lung cells and promote activation of integrated HIV proviruses that, making the lung epithelial cells as an important reservoir of HIV. Therefore, it is feasible that emerging respiratory infections, like current COVID19 (coronavirus-induced disease of 2019) pandemic may complicate the lung disease phenotypes among PLWH (40–42). And the ongoing immunosuppression and antiretroviral-mediated altered immune responses could further result in severe outcomes.

## MATERIALS & METHODS

### Normal Human Bronchial Epithelial Cells and ALI culture

Normal Human Bronchial Epithelial Cells (NHBE) cells were obtained from MatTek Incorp (EpiAirway^™^, Ashland, MA). For all air-liquid interface (ALI) cultures of primary NHBEs (Lonza Inc., Basel, Switzerland), cells were plated onto collagen IV-coated 24-mm Transwell-clear culture inserts (Corning Costar Corporation, Cambridge, MA) at a density of 5×10^5^ cells/cm^2^ in BEGM media (Lonza Inc) and subsequently in ALI media as described previously (43). The apical surface of the cells was exposed to air and cells were cultured for another 21 days till they were fully differentiated.

### HIV Infection and p24 Analysis

The X4-tropic viral strain HIV-1_LAV_ and R5-tropic viral strain HIV-1BaL were employed in these studies. NHBE ALI cultures grown on transwells were infected apically and basolaterally with of either X4-tropic HIV-1_LAV_ or R5-tropic HIV-1_BaL_ (5 ng/ml p24 equivalent) as described earlier (44). 16 hours post infection, cells were washed apically and basolaterally with PBS four-times to remove any residual input virus. The fourth wash was collected for p24 analysis and measured as day 0 to confirm that all input virus had been removed. Culture supernatants were collected on day 3 and p24 antigen levels were determined using p24 ELISA (ZeptoMetrix Corp. Cat # 0801200) as per manufacturer’s instructions. For chronic HIV exposure, NHBE ALI cultures were infected with X4-tropic HIV strain (IIIB) or R5-tropic HIV strain (BaL) (2.5 ng/ml p24 equivalent) were allowed to proceed for 7 days.

### Immunostaining and Fluorescent Imaging Analysis

For immunohistochemical staining, deparaffinized and hydrated lung tissue sections were washed in 0.05% v Brij-35 in PBS (pH 7.4) and immunostained for antigen expression as described previously (19). Briefly, the antigens were unmasked by steaming the sections in 10 mM Citrate buffer (pH 6.0) followed by incubation in a blocking solution containing 3% BSA, 1% Gelatin and 1% normal donkey serum with 0.1% Triton X-100 and 0.1% Saponin and were stained with antibodies to CD4 (Abcam, #ab133616), CXCR4 (Abcam, #ab181020), CCR5 (Abcam, #ab11466), HIV-Tat (Abcam, #ab63957) and pan-CK (#4545, Cell Signaling Technologies, Danvers, MA). The NHBE cells grown on coated coverslips or the coated Labtek-II slides (Thermo Fisher Inc.) were fixed in 4% paraformaldehyde and washed in 0.05% v Brij-35 in PBS (pH 7.4) and immunostaining was performed as described previously (45). Briefly, the cells were blocked using a solution containing 3% BSA, 1% Gelatin and 1% normal donkey serum with 0.1% Triton X-100 and 0.1% Saponin and were stained CD4, CXCR4, CCR5 and pan-CK, as described above. The immunolabelled cells/tissue sections were detected using respective secondary antibodies conjugated fluorescent dyes (Jackson ImmunoResearch Lab Inc., West Grove, PA) and mounted with 4’,6-diamidino-2-phenylindole (DAPI) containing Fluormount-GTM (SouthernBiotech, Birmingham, AL) for nuclear staining. Immunofluorescent images were captured using BZX700 Microscopy system (Keyence Corp., Japan) and analyzed using NIH Image J software.

### Quantitative Real-Time RT-PCR

Total RNA was isolated from the experimental cells using RNAeasy kit (Qiagen, Germantown, MD) as per manufacturer’s instruction. RNA concentration was determined using the Synergy HTX Multi-Mode reader (BioTek, Winooski, VT) and cDNA were synthesized using iScript advanced cDNA kit (BioRad, Hercules, CA). The primer/probe sets for CD4, HIV-LTR RNA, and MUC5AC were obtained from Applied Biosystems (Thermo Fisher Inc.) and cDNA amplified was quantified by q-PCR using the TaqMan Gene expression kit (Thermo Fisher Inc.) in the Agilent Stratagene Mx3000P Real-Time PCR System (Thermo Fisher Scientific, Waltham, MA). Relative quantities were calculated by normalizing averaged CT values to CDKN1B or GAPDH to obtain ΔCT, and the fold-change (ΔΔCT) over the controls were determined as described previously (45).

### Western Blot Analysis

Cell extracts were prepared using RIPA buffer (20 mM Tris, pH 7.4, 137 mM NaCl, 1% NP-40, 0.25% Deoxycholate, 0.1% SDS, 1 mM EDTA and 1% protease inhibitor cocktail). Protein concentration was determined by BCA kit (Pierce; Rockford, IL) and 50 μg protein was analyzed by western blotting as described previously (43). Antibodies used were for CCR5 (Abcam, #ab11466) and for β-actin (Sigma Co. St. Louis, MO). Proteins were detected using ECL and visualized by chemiluminescence (Perkin Elmer, Waltham, MA) using the BioRad Chemidoc Imaging system (Hercules, CA).

### Alu-PCR and Nucleotide Sequencing

We utilized the methods as described previously (46) in which two-step Alu-gag PCR assay was performed to confirm the presence of integrated HIV-1 proviral DNA. This method utilizes a nested PCR approach. Briefly, the during 1^st^ round PCR, we amplified a region between HIV-1 gag gene and nearest Alu repeat element of host genome. The primer sequences used in the 1st step PCR were: Alu Forward: 5’-GCCTCCCAAAGTGCTGGGATTACAG - 3’ and HIV gag reverse: 5’-GTTCCTGCTATGTCACTTCC - 3’ (corresponds to 1505-1486 of HXB2 genome). The reactions were carried out in 50 μl containing 1.5 mM MgCl2, 0.2 mM dNTPs mix, 100 nM of Alu Forward primer, 600 nM of HIV gag reverse primer and 5 U of Platinum Taq DNA polymerase (Life Technologies; USA). In the 2^nd^ round PCR 5’ LTR region of HIV-1 was amplified using 1 μl of 1^st^ round PCR amplicon as a template. The primers used for second round were: 5’ LTR Forward: 5’-TTAAGCCTCAATAAAGCTTGCC - 3’ and 5’ LTR Reverse: 5’-GTTCCTGCTATGTCACTTCC - 3’.). The reactions were carried out in 50 μl containing 1.5 mM MgCl2, 0.2 mM dNTPs mix, 400 nM of 5’ LTR Forward primer, 400 nM of 5’ LTR Reverse primer and 5 U of Platinum Taq DNA polymerase. The PCR amplification was carried out in an Applied Biosystem 9700 thermal cycler with the following program with following conditions, 1st Round PCR: 2 min at 95°C, followed by 40 cycles of denaturation at 95°C for 15s, annealing at 50°C for 15s and extension at 72°C for 3 min 30s; 2^nd^ round PCR: 2 min at 95°C, followed by 40 cycles of denaturation at 95°C for 15s, annealing at 60°C for 15s and extension at 72°C for 30s. Finally, PCR amplicons were resolved on a 2% agarose gel (Promega Corporation, Madison, USA) pre-stained with Ethidium bromide (0.5 μg/ml) and gel images were documented using a Bio Rad gel documentation system (BIORAD, USA). Next, the PCR amplicons were purified using QIAGEN PCR purification kit (QIAGEN, Germany) as per the manufacturer’s instructions and subjected to Sanger sequencing using BigDye Terminator v3.1 Cycle Sequencing Kit (Applied Biosystems, California; USA). Automated capillary electrophoresis was performed on an ABI PRISM^®^ 3500 Genetic Analyzer (Applied Biosystems, California; USA) using data collection software v.3.1 at the UNMC DNA Sequencing Core facility. Raw sequence data were manually edited, spliced and assembled by Sequencher v4.9 to generate the final contig. Pairwise sequence alignment of edited sequence was performed with HXB2 reference sequence of HIV-1 using Clustal W (47).

### RNA Fluorescent In-Situ Hybridization (FISH)

RNA FISH was essentially performed using the RNAscope^®^ Fluorescent Multiplexed reagent kit (Advanced Cell Diagnostics, Newark, CA) as per the manufacturer’s protocol. The probe set for HIVgag-pol consisted of 20 dual probes targeting different segments within the whole transcript (Advanced Cell Diagnostics). Formalin-fixed paraffin-embedded airway sections were deparaffinized and permeabilized using 0.1% PBS Triton-X100 for 10 min at RT followed by washing in PBS. Probes were hybridized for 2 h at 40°C using a HyBEZ^®^ oven (Advanced Cell Diagnostics). The signal was amplified by subsequent incubation of Amp-1, Amp-2, Amp-3 and HRP-tagged probe (Thermo Fisher Inc), one drop each for 30, 15, 30, and 15 min, respectively, at 40°C using a HyBEZ^®^ oven. Each incubation step was followed by two-times 2 min washes using RNAscope washing buffer in slide holders with agitation (50 rpm). Glass slides were applied into slide holder containing washing buffer. The probes were then detected using Tyramide signal amplification (TSA) reaction using an Alexa-flour labelled TSA kit (PerkinElmer Bioscience) based on the manufacturer’s instructions. The sections were processed for immunostaining of Pan-CK (--) as described above with 4’,6-diamidino-2-phenylindole (DAPI) containing Fluormount-G (SouthernBiotech, Birmingham, AL) to visualize nuclei.

Immunofluorescent images were captured with BZX700 Microscopy system (Keyence Corp, Japan) and analyzed by NIH ImageJ software. RNA FISH expressions were quantified by RNAscope data analysis suggested by Advanced Cell Diagnostics, (Newark, CA, USA). Each probe signals (dots)/cell were counted and allotted for separate bins as follows: Bin 0 (0 Dots/Cell); Bin 1 (1-3 Dots/Cell); Bin 2 (4-9 Dots/Cell); Bin 3 (10-15 Dots/Cell); Bin 4 (>15 Dots/Cell). Then, the Histo score (H-Score) was calculated as follows: H-Score = Sum of each (bin number x percentage of cells per bin) i.e., H-score = (0 × % cells in Bin 0) + (1 × % cells in Bin 1)+(2 × % cells in Bin 2) + (3 × % cells in Bin 3) + (4 × % cells in Bin 4). Final scores derived by this metric will have a range between 0 and 400.

### Statistical Analysis

Grouped results were expressed as means ± SEM. Data were analyzed using GraphPad Prism Software (GraphPad Software, Inc., San Diego, CA). Grouped results were analyzed using two-way analysis of variance. When significant main effects were detected (P < 0.05), Fishers least significant difference test was used to determine differences between groups.

## ACKNOWLEDGEMENTS

Authors appreciate the technical support provided by Ruben Castro. Authors acknowledge the funding support by NIH R01HL125000 (to M.S.), and R21AI144374, R21AI117560, R01 HL147715, and the FIU Start-Up Funds (to H.S.C.).

