## Supplementary material for "Lung Bronchial Epithelial Cells are HIV Targets for Proviral Genomic Integration": Online Supplemental Data

[#] Equal Contributions

**Running Title**: Bronchial Epithelial Cells are HIV Targets

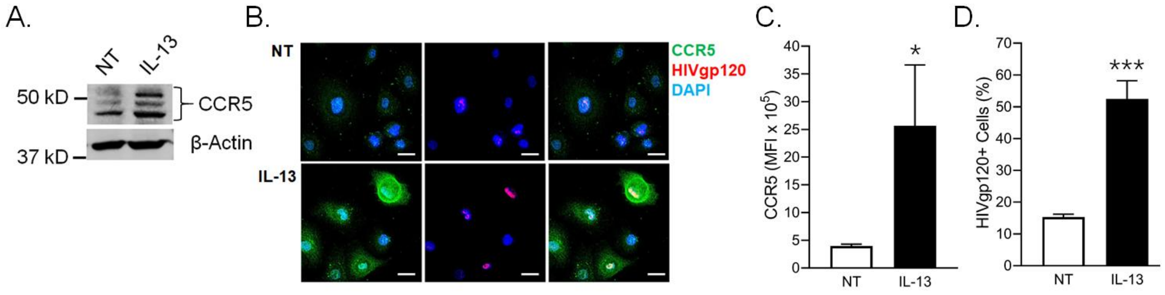


**Figure S1.** Induced expression of CCR5 following IL-13 treatment of NHBE cells. (**A.**) Western blot analysis of CCR5 isoforms expression in IL-13 treated NHBE cells compared to non-treated (NT) cells, β-actin levels used as internal controls. (**B.**) Micrographs of NT and IL-13 treated NHBE cells showing CCR5 (green) expression and immunopositivity for HIVgp120 (red) along with DAPI-stained nuclei (blue); scale - 10μ. (**C.**) Mean Fluorescence Intensity (MFI) of CCR5 expression and (**D.**) quantification of HIVgp120+ cells for each treatment. Data shown as mean±SEM; n=3/gp; *p< 0.05, ***p<0.001.

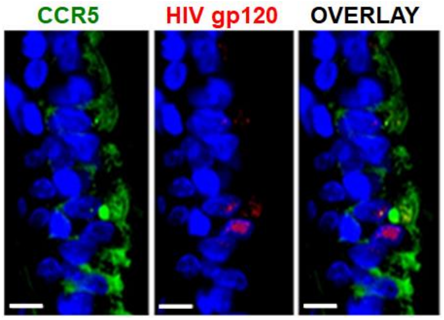


**Figure S2. CCR5 expression in the lung epithelial cells of SHIV-infected nonhuman primates.** Representative micrograph of CM bronchial epithelial cells showing CCR5 (green) expression in CS+SHIV-infected CMs and co-detection of HIV gp120 (red) along with the DAPI-stained nuclei (blue); scale - 5µ.

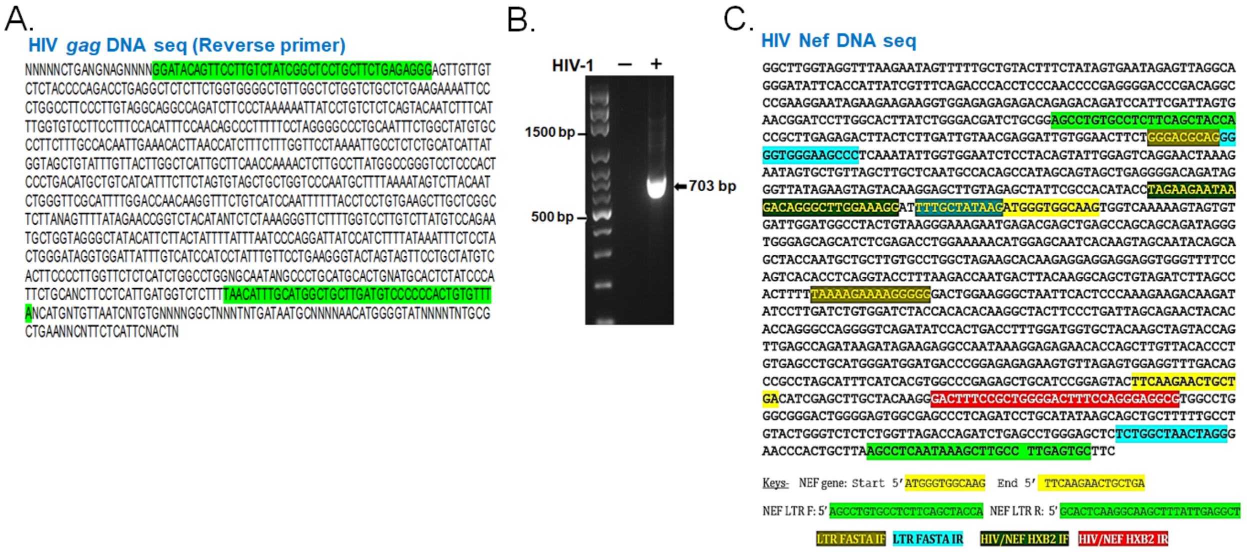


**Figure S3. Amplification and sequencing analysis of HIV gag and Nef DNA to confirm integration of HIV-1 proviral genome in NHBEs. (A.)** HIV gag DNA sequence obtained using the reverse primers that are highlighted in green shade. **(B.)** Agarose gel analysis of the HIV Nef PCR amplicon (703 bp) amplified from the DNA isolated from HIV-infected NHBEs. **(C.)** HIV Nef DNA sequence obtained from sequencing of the 703 bp PCR product. The primer sequences used are highlighted in green and 5'-LTR sequence of HIV is highlighted in blue and other identified sequences are indicated in the key below.
